# The heritability and structural correlates of brain entropy

**DOI:** 10.1101/2024.03.07.583865

**Authors:** Yi Zhen, Yaqian Yang, Yi Zheng, Xin Wang, Longzhao Liu, Zhiming Zheng, Hongwei Zheng, Shaoting Tang

**Author notes:** Email addresses:* (Hongwei Zheng), (Shaoting Tang).

## Abstract

Brain entropy (BEN) quantifies the temporal complexity of spontaneous neural activity, which has been increasingly recognized as providing important insights into cognitive functions and psychiatric disorders. However, its heritability and structural underpinnings are not well understood. Here, we utilize sample entropy to calculate BEN in healthy young adults using resting-state fMRI from the Human Connectome Project. We show that BEN is significantly heritable in broad brain regions. Heritability estimates exhibit regional variability, with lower heritability in the limbic and subcortical networks than in other networks. We further relate BEN to structural features including surface area (SA), cortical myelination (CM), cortical thickness (CT), and subcortical volumes (SV). We find that BEN is negatively correlated with SA in most cortical regions, but is positively correlated with CM in the prefrontal cortex, lateral temporal lobe, precuneus, lateral parietal cortex, and cingulate cortex. Only a few cortical regions show significant correlations between BEN and CT. By decomposing correlations into genetic and environmental components, we observe that most of BEN-SA correlations are associated with shared genetic and environmental effects. BEN-CM correlations are mainly attributed to shared environmental effects. BEN-CT correlations are partly related to shared environmental effects. We find no significant associations between BEN and SV. Collectively, our work establishes the genetic basis and structural correlates of resting-state fMRI-based BEN, supporting its potential application as an endophenotype for psychiatric disorders.

## 1. Introduction

The human brain is a dynamic and complex system that exhibits ongoing fluctuations in neural activity even in the absence of explicit tasks [1, 2, 3]. The temporal dynamics of brain activity provide a scaffold for flexible information processing that supports sophisticated cognitive behaviors [4, 5, 6]. Dissecting these temporal dynamics is instrumental for understanding cognitive functions and pathological substrates of mental diseases. Temporal complexity analysis using the sample entropy of fMRI signal [7, 8, 9, 10], often referred to as brain entropy (BEN), is an emerging method for tracking the temporal dynamic properties of neural activity. Sample entropy is a well-defined measure derived from information theory that quantifies the predictability, irregularity, and complexity of a signal [11]. Larger entropy values indicate that a signal is less predictable and more complex, while smaller entropy values mean that a signal is more predictable and less complex [11, 12]. In comparison to other types of entropy measure (such as Shannon entropy [13] and approximate entropy [14]), sample entropy is less sensitive to noise and allows for more accurate entropy estimation for signals with a limited number of sampling points, rendering it suitable for the complexity analysis of fMRI [11, 15, 7]. Previous studies have demonstrated that BEN can accurately discriminate between empirical fMRI and simulated signals [7] and show good reproducibility [16, 17] and task-induced sensitivity [9, 18, 19]. Evidence from theoretical models, animal experiments, and imaging data analyses suggests that BEN may inform the capacity of brain information processing at local and distributed levels [20, 21, 22]. Moreover, this entropy measure is theoretically independent of commonly used variance-based temporal measures such as signal variance or standard deviation, and thus may provide a new avenue for understanding brain dynamics and functions.

Previous studies have demonstrated that BEN is sensitive to disease-induced brain alterations. Aberrant BEN has been found in a host of neuropsychiatric diseases including Alzheimer’s disease [23, 24], schizophrenia [25], autism spectrum disorder [26], attention deficit hyperactivity disorder [27], major depressive disorder [28, 29] and generalized anxiety disorder [30]. Notably, one previous fMRI study used multiscale sample entropy to study mild Alzheimer’s disease [31]. They found that the mild Alzheimer’s disease group had significantly lower entropy than the control group in default mode regions whereas there were no significant differences in functional connectivity between the groups, implying that multiscale BEN might be more sensitive in identifying cognitive decline relative to functional connectivity. Recently, there has been a growing interest in investigating individual-specific complexity profiles. Several studies have reported remarkable inter-individual variability in BEN [17, 32]. The individual BEN profiles can serve as a “fingerprint” to accurately distinguish subjects from a large group [17], and are related to task activation [33] and individual behavioral traits [17, 32, 34]. These findings suggest that BEN may act as biomarkers for the diagnosis of brain disorders and provide valuable insight into individual differences in cognitive function.

By combining neuroimaging and twin information, numerous studies have demonstrated that brain structural characteristics such as cortical thickness [35], myelination [36], cortical folding [37, 38], and white-matter microstructure [39, 40] are heritable. Focusing on measures derived from fMRI data, prior studies have shown that functional interaction features between brain regions such as effective connectivity [41], and static and dynamic functional connectivity [42, 43, 44, 45] are heritable. Heritable imaging measures hold promise as intermediate phenotypes for exploring the pathogenesis of brain disorders [43, 46]. However, to date it remains unclear whether there exists a genetic underpinning for BEN. In view of the potential cognitive and clinical implications of BEN, it is necessary to disentangle the degree to which genetic factors contribute to BEN.

The relationship between structure and function is a fundamental question in neuroscience. With network neuroscience methods, numerous investigations have found that white matter fiber pathways can to some extent account for inter-regional functional interactions [47, 48, 49, 50, 51]. Previous work has reported that the amplitude of low-frequency fMRI fluctuations is correlated with the brain volume [52]. Inspired by these observations, we conjecture that BEN may also be somewhat constrained by brain structure. Recent studies have provided preliminary evidence that the temporal complexity of fMRI is associated with white matter integrity [53], cortical folding [54], and myelination [54]. However, these findings only focused on the phenotypic correlation level, lacking an understanding of the genetic and environmental overlap between structure and complexity. Also, these findings are generally based on the coarse-scale network level or limited to cortical areas, lacking a comprehensive description of fine-scale regional structure-complexity coupling on the whole brain containing both cortical and subcortical areas.

In this study, our aim is to investigate whether BEN is controlled by genetic effects and related to brain structure. We first employ a classical twin design in a large adult sample to estimate the heritability of BEN. Next, we utilize multiple linear models to assess phenotypic associations between BEN and brain structure including cortical surface area (SA), cortical myelination (CM), cortical thickness (CT), and subcortical volumes (SV). Finally, we conduct bivariate genetic analyses to evaluate whether shared genetic or environmental factors drive phenotypic associations.

## 2. Materials and methods

### 2.1. Participants

In this study, we used the HCP S1200 dataset [55], which consists of 1206 healthy adult subjects. Subjects without four complete resting-state fMRI scans were removed. We prioritized the use of genotyping information to confirm each participant’s zygosity information when possible, otherwise we used self-reported zygosity information. We excluded three participants who lacked zygosity information. Finally, a total of 1001 participants (533 women and 468 men) aged between 22 and 37 were used for subsequent analyses, which included 132 monozygotic twin pairs, 64 dizygotic twin pairs, 101 non-twin siblings, and 130 singletons. The HCP project was approved by the Institutional Review Board at Washington University in St. Louis, and all participants provided written informed consents.

### 2.2. Image Acquisition and Processing

The HCP MRI data were acquired on a customized Siemens 3T Connectome Skyra scanner with a 32-channel RF-receive head coil at Washington University in St. Louis. Each subject had four 14.4-minute resting-state fMRI runs and each run contained 1200 volumes. The detailed setting of HCP acquisition can be found in [56]. All functional acquisitions were preprocessed according to the HCP minimal preprocessing pipeline [57], and detailed steps included echo-planar imaging gradient distortion, motion correction, image distortion correction, spatial transformation to the Montreal Neurological Institute (MNI) space, and intensity normalization. ICA-FIX was used to remove the artifact [58, 59]. In addition, accurate inter-subject registration of cortical surfaces was performed using the Multimodal Surface Matching algorithm (“MSMAII”) [60, 61]. In the robustness analysis, we additionally performed global signal regression (GSR) and evaluated its impact on our results. We parcellated the cortical gray matter into 400 regions using the Schaefer-400 atlas [62]. 19 subcortical regions from FreeSurfer subcortical segmentation were also included in this study [63]. Structural images were preprocessed by the minimal preprocessing pipeline mentioned above. Cortical thickness (CT) and cortical surface area (SA) were estimated through surface reconstruction from FreeSurfer [64]. Cortical myelin (CM) content was estimated using the ratio of T1w/T2w image intensities [65]. The Schaefer-400 atlas was also used for structure images to obtain regional cortical measures. The regional surface area was measured as the sum of the areas of all vertices within the region. The regional cortical thickness was estimated by averaging the thickness of all vertices within the region. The regional cortical myelination was calculated by averaging the myelination of all vertices within the region. The volumes of 19 subcortical regions were obtained from FreeSurfer automatic segmentation [63].

### 2.3. The calculation of BEN

Sample entropy was used to estimate the BEN of regional average fMRI time series for each of four preprocessed runs, resulting in four BEN maps for each subject. The four maps were further averaged together, and the resulting mean BEN map for each subject was used for subsequent analyses. A brief description of how we calculated this entropy measure is as follows.

The basic calculation steps of sample entropy are in line with [11, 17, 66]. Given an input signal (such as an fMRI time series), *X* = (*x*_1_, *x*_2_, *x*_3_, *· · ·, x*_*N*_), we first constructed an m-dimensional embedding vector,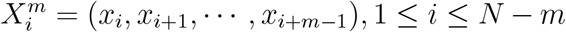. Then for each *i*, we defined

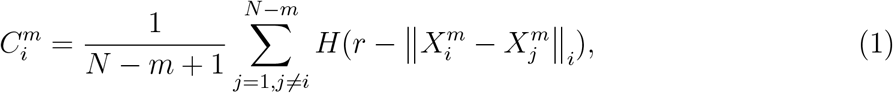

where *r* is tolerance parameter, *H*(*•*) represents the Heaviside function, and ∥•∥_1_ denotes the Chebyshev distance. Next, for each *i*, we defined

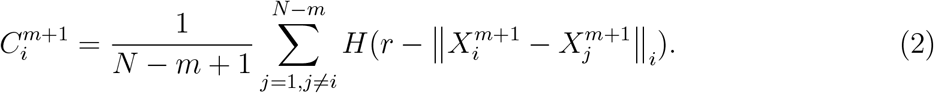

Finally, the sample entropy of *X* was given by:

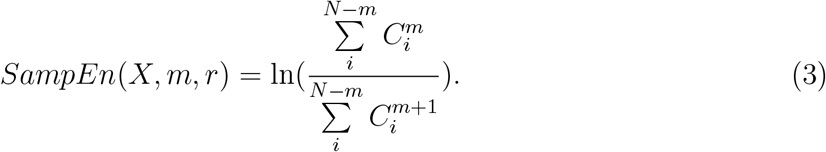

Consistent with the parameter settings of extensive literature, we chose *m* = 2 and *r* = 0.5*σ*_*X*_ for BEN calculation. It has been demonstrated that *m* = 2 and *r* = 0.5*σ*_*X*_ can provide highly reproducible for the complexity analysis of fMRI signals [16]. In the robustness analysis, we additionally included another pair of widely used parameters (*m* = 3 and *r* = 0.6*σ*_*X*_) and assessed the influence of different parameter settings on our findings.

### 2.4. Statistical Analyses

Phenotypic correlation analyses between BEN and brain structure were performed at the regional level using multiple linear regression models. More specifically, SA, CM, and CT for each cortical region, as well as SV for each subcortical region, were regressed on the corresponding regional BEN respectively, while controlling for age, sex, age^2^, age*×*sex, age^2^*×*sex, and total intracranial volume. The results of phenotypic analyses were corrected by the Bonferroni method, and the significance threshold was set to *p <* 0.05*/*(400 *×* 3 + 19).

The heritability of regional BEN and brain structure were estimated using the R package solarius [67], which was an R interface to the Sequential Oligogenic Linkage Analysis Routines (SOLAR) software. SOLAR employed a maximum likelihood variance decomposition algorithm to divide phenotypic variance into an additive genetic component and an environmental component [68, 69]. Narrow-sense heritability (*h*^2^) was represented as the proportion of the variance in a phenotype 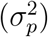 attributable to the additive genetic component 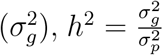. To determine if common genetic or environmental factors concurrently influence BEN and brain structure, genetic and environmental correlations between BEN and brain structure were estimated by bivariate genetic analyses. SOLAR implemented bivariate genetic analyses based on the following formula:

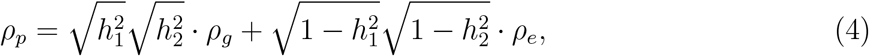

where the phenotypic correlation (*ρ*_*p*_) was partitioned into the genetic correlation (*ρ*_*g*_) and the environmental correlation (*ρ*_*e*_). 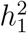 and 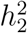 represented the heritability estimates for BEN and brain structure [70, 43, 71, 72]. If *ρ*_*g*_ is significantly different from zero, a genetic overlap (pleiotropy) between two phenotypes is considered [70, 43]. In this study, all heritability analyses controlled for demographic covariates including age, sex, age^2^, age*×*sex, and age^2^*×*sex. In addition, when investigating brain structure heritability, and genetic and environmental correlations, total intracranial volume was also included as a covariate, which was consistent with phenotypic correlation analyses. Prior to all genetic analyses, inverse normal transformations were performed to ensure the normality of all imaging phenotypes [43]. For the univariate heritability analysis, results were corrected by the Bonferroni method, which was performed separately for BEN and brain structure. For the bivariate genetic analysis, results were corrected by the FDR method (corrected *p <* 0.05) [73], which was performed separately for genetic and environmental correlation analyses.

## 3. Results

### 3.1. Heritability of BEN

We performed heritability analyses of regional BEN, with covarying for age, sex, age^2^, age*×*sex, age^2^*×*sex. Results of heritability estimates were shown in Figure.1. We found that BEN was significantly heritable in most brain regions (398/419 regions) and heritability estimates were moderate, with the heritability ranging from 0 to 0.68 (*h*^2^ : *M* = 0.458, *SD* = 0.130). High heritability estimates were found in the occipital lobe, fusiform cortex, precuneus, lateral parietal cortex, lateral temporal cortex, middle frontal cortex, and isthmus of the cingulate cortex, while low estimates were observed in the parahippocampal cortex, orbitofrontal cortex, cingulate, entorhinal cortex, temporal pole, bilateral nucleus accumbens, left ventral diencephalon, and bilateral pallidum. We noted that there was a strong negative correlation between the magnitude of BEN and its heritability (*r* = *−*0.58, *p <* 0.0001), indicating that higher BEN was related to weaker genetic effects. We further grouped heritability estimates into eight functional networks [74], including the visual (VIS), somatomotor (SM), limbic (LIM), dorsal attention (DAN), ventral attention (VAN), control (CON), default mode (DMN), and subcortical (SUB) networks. The heritability of BEN in LIM and SUB was significantly lower than in other networks (corrected *p <* 0.05, Bonferroni correction).

**Figure 1.**
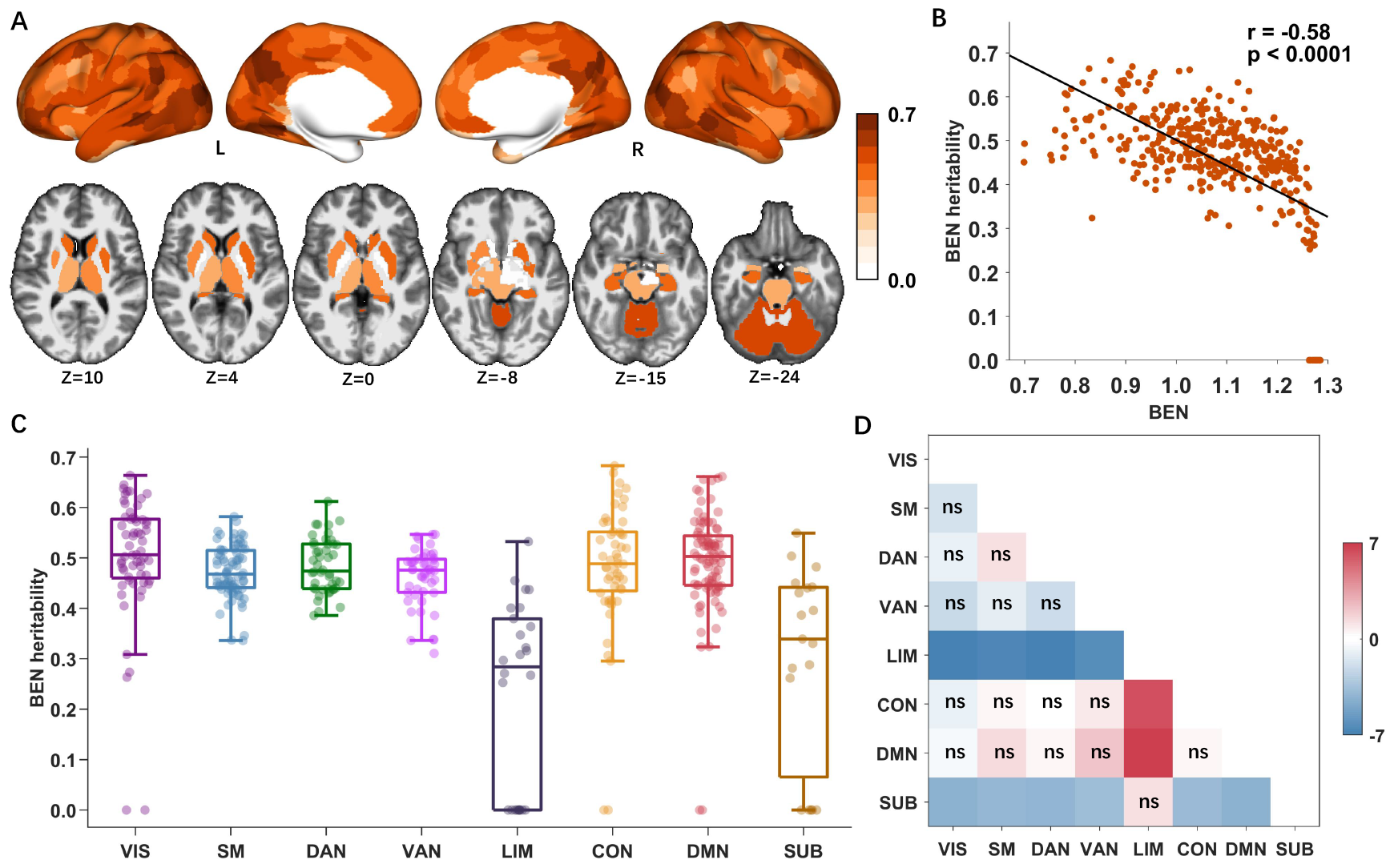
Regional BEN heritability estimates. A. Heritability map of regional BEN. B. Spatial correlation between BEN heritability estimates and BEN magnitude (Pearson *r* = *−*0.58, uncorrected *p <* 0.0001). C. Heritability estimates of regional BEN grouped by 8 brain networks: visual (VIS), somatomotor (SM), dorsal attention (DAN), ventral attention (VAN), limbic (LIM), control (CON), default mode (DMN), subcortical (SUB). Boxplots are combined with scatter plots, where each point denotes the heritability estimate of a brain region, the central line of each box indicates the median, the top/bottom edge of each box denotes the 75th/25th percentiles, and the whiskers of each box indicate the upper/lower bounds of 1.5*×*the interquartile range. (D) The t-statistics for pairwise comparisons between brain networks. Each pairwise comparison was performed using the two-sided t-test and computed as the brain network on the bottom axis relative to the brain network on the right axis. All p values were corrected using the Bonferroni correction (significance threshold at *p <* 0.05*/*28), and non-significant comparisons were labeled as ns. The heritability estimates of regional BEN in LIM and SUB were significantly lower than other networks.

### 3.2. Phenotypic correlations between BEN and brain structure

We performed phenotypic correlation analyses between cortical structure (SA, CM, and CT) and BEN for each cortical region, as well as between SV and BEN for each subcortical region. All analyses controlled the effects of age, sex, age^2^, age*×*sex, age^2^*×*sex, and intracranial volume. As shown in Figure.2, significant negative correlations between SA and BEN were observed in most regions of the cortex. Several positive correlations were observed in the paracentral lobule and right lingual gyrus. Regions with no significant associations were primarily located within the medial visual and sensorimotor cortices. Phenotypic associations between BEN and CM were in the opposite direction to those widely observed with SA. Significant positive correlations emerged in multiple cortical regions, encompassing the prefrontal cortex, lateral temporal lobe, precuneus, lateral parietal cortex, and cingulate cortex. No significant negative correlations were detected. These results indicated that higher local CM was associated with larger entropy and complexity. In contrast to SA and CM, significant phenotypic associations between CT and BEN were observed in relatively few cortical regions. Negative correlations were found in the frontal lobe, parietal lobe, insula, right lateral temporal cortex, and cingulate cortex. Positive correlations were found in the orbitofrontal gyrus, right cingulate/precuneus cortex, left sensorimotor cortex, and right visual cortex. We didn’t observe significant phenotypic relationships between SV and BEN.

**Figure 2.**
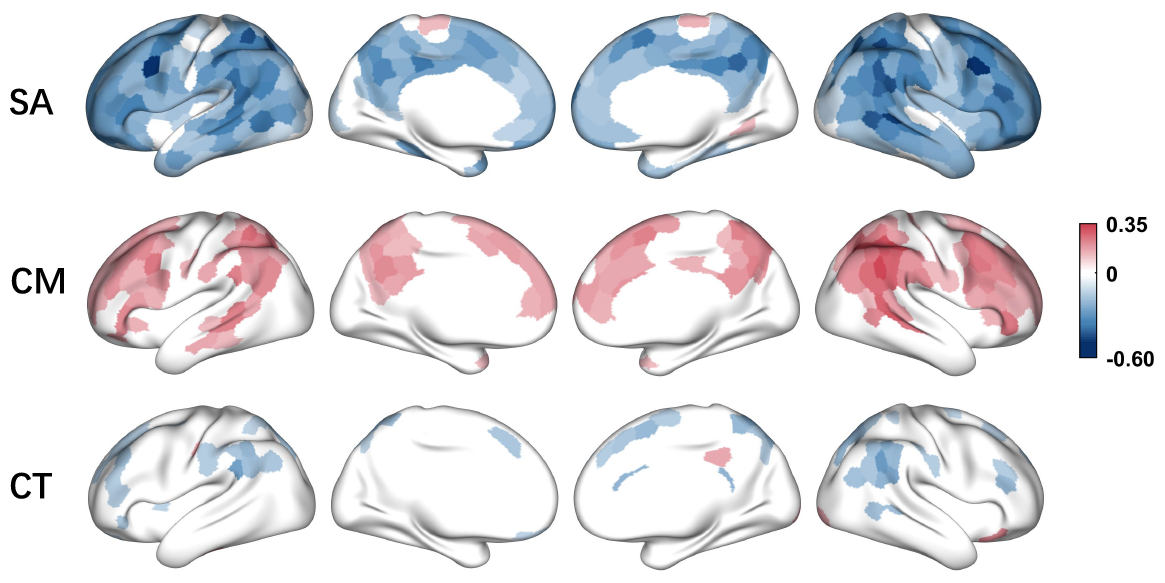
The phenotypic correlations between cortical structure (surface area (SA), cortical myelination (CM), cortical thickness (CT)) and regional BEN. All results are corrected using the Bonferroni method (significance threshold at *p <* 0.05*/*1219)

### 3.3. Genetic and environmental correlations between BEN and brain structure

We further assessed whether phenotypic correlations between BEN and brain structure were driven by common genetic and environmental factors. We first evaluated the heritability of local brain structure (Figure.3). We observed that regional SA (*h*^2^ : *M* = 0.526, *SD* = 0.106), CM (*h*^2^ : *M* = 0.482, *SD* = 0.096), and CT (*h*^2^ : *M* = 0.390, *SD* = 0.120) were heritable in most cortical regions, and all subcortical volumes were heritability (*h*^2^ : *M* = 0.685, *SD* = 0.115). Heritability maps of brain structure were highly consistent with previous findings [36, 37, 72].

**Figure 3.**
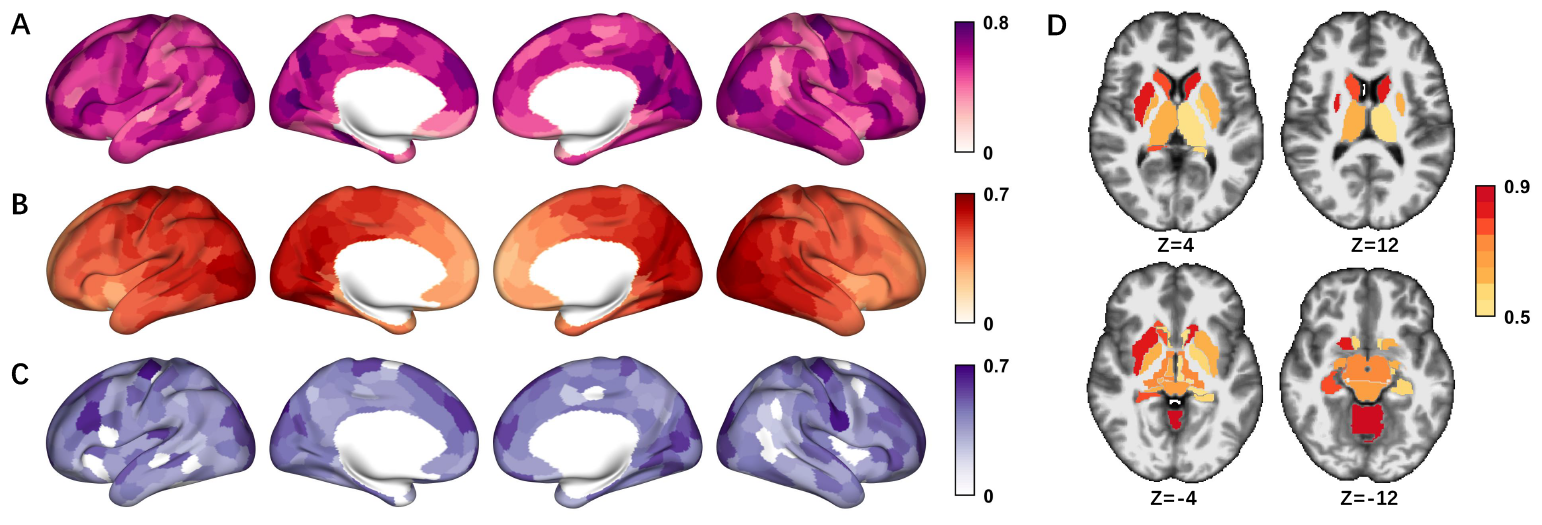
Local brain structure heritability estimates. A. Heritability map of cortical surface area. B. Heritability map of cortical myelination. C. Heritability map of cortical thickness. D. Heritability map of subcortical volumes. All results are corrected using the Bonferroni method (significance threshold at *p <* 0.05*/*1219)

Subsequently, we performed region-wise bivariate heritability analyses to examine genetic and environmental correlations between BEN and brain structure. The regional structure-entropy genetic and environmental correlation maps were shown in Figure.4. Bivariate heritability analyses showed that most of the phenotypic associations between SA and BEN were driven by shared genetic and environmental effects. Negative genetic and environmental correlations were observed in most cortical regions. One positive environmental correlation was found in a small region of the paracentral lobule. There was little evidence to support that the phenotypic associations between CM and BEN were primarily attributable to common genetic factors, with only a few significant genetic correlations in the lateral temporal lobe, parietal lobe, cingulate, and prefrontal cortex. Conversely, the majority of phenotypic associations were associated with significant environmental correlations. More specifically, positive environmental correlations were mainly observed bilaterally in the prefrontal cortex, lateral temporal cortex, cingulate, and parietal cortex. Sparse phenotypic associations between CT and BEN that we observed were, in part, driven by shared environmental factors. Significant shared genetic effects were barely detected, with only phenotypic correlations in two small regions of the right superior frontal cortex related to genetic correlations.

**Figure 4.**
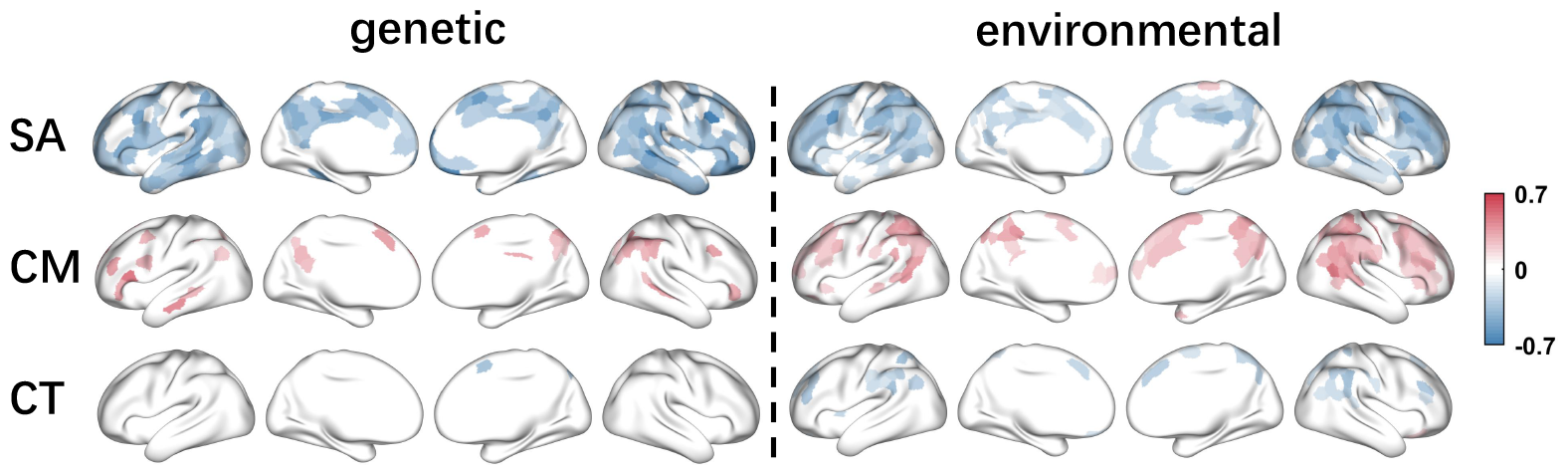
The genetic and environmental correlations between cortical structure (surface area (SA), cortical myelination (CM), cortical thickness (CT)) and regional BEN. All results are corrected via the FDR correction (FDR corrected *p <* 0.05). Results only show regions with significant regional BEN heritability, significant phenotypic correlation, and significant cortical structure heritability.

### 3.4. Validation analyses

We performed two additional analyses to verify the robustness of our findings. Firstly, we repeated our findings in another frequently adopted parameter setting. We found that two different parameter choices yielded highly consistent results in heritability estimates (Pearson *r* = 0.99) and structure-entropy relationships (see Supplementary Fig.S1). Secondly, we additionally included GSR in the fMRI preprocessing process. We observed that the analyses with GSR were highly similar to the analyses without GSR in heritability estimates (Pearson *r* = 0.91) and structure-entropy relationships (see Supplementary Fig.S2), indicating that GSR didn’t affect our findings.

## 4. Discussion

In this paper, we focused on the heritability and structure correlates of resting-state fMRI-based BEN. Using the twin and imaging information from the HCP database, we provided robust evidence that regional BEN was significantly heritable in most brain regions (398/419 regions). The magnitude of heritability was variable across the brain, with limbic and subcortical networks exhibiting lower heritability estimates relative to other networks. We further explored the relationships between BEN and local brain structure (SA, CM, CT, and SV). For SA, we observed negative phenotypic correlations in widespread cortical regions (except for median visual and somatomotor regions), and that most of these correlations were associated with genetic and environmental correlations. For CM, we observed positive phenotypic associations in the prefrontal cortex, lateral temporal lobe, precuneus, lateral parietal cortex, and cingulate cortex, with these phenotypic associations mainly attributed to shared environmental effects. For CT, we observed sparse correlations, which were partly associated with environmental correlations. For SV, no significant association was observed. Collectively, these results offer insights into the genetic basis and structural substrates of BEN, promoting its further applications in understanding cognitive functions and neurological disorders.

### 4.1. Heritability of BEN

The brain is a complex system whose functionality emerges from neural activities with ongoing fluctuations. BEN is an emerging approach in the neuroimaging field, which has provided important perspectives for the temporal dynamics of brain signal fluctuations. However, its genetic basis remains poorly explored, which limits insights into the biological underpinnings of brain dynamics and hinders further clinical biomarker applications. Here, through the heritability analysis, we demonstrated that BEN was genetically controlled. More specifically, we observed moderate heritability, with an average of about 46% of the inter-individual variance attributed to additive genetic effects. The heritability estimates are comparable to other fMRI-derived temporal dynamic characteristics, such as the functional network activity represented by the variance of network temporal fluctuations (*h*^2^ : 0.23 *−* 0.65) [75], the variance of dynamic functional connectivity (*h*^2^ : 0.2 *−* 0.59, *mean* = 0.45) [44], and dynamic features of connectome configuration transitions (fractional occupancy: *h*^2^ = 0.39, transition probability: *h*^2^ = 0.43) [45], slightly lower than complexity measures of EEG signals such as long-range temporal correlations quantified via Hurst exponent (*h*^2^ : 0.33 *−* 0.6) [76] and the pointwise correlation dimension (*h*^2^: eyes-closed condition: 0.62-0.68; eyes-open condition: 0.46-0.6) [77]. Our findings, combined with these previous studies, provide converging evidence that genetic factors play an indispensable role in shaping brain temporal dynamics. Besides, we showed that the heritability of regional BEN varied across brain regions. We observed a significant negative association between BEN value and its heritability, i.e., low complexity tended to show high heritability, suggesting that genetic effects appear to play a preferential role in sculpting more predictable (lower entropy) brain activities. We also found that limbic and subcortical networks exhibited significantly lower heritability than other regions. The low signal-to-noise ratio of fMRI signals in limbic and subcortical regions could be one possibility driving the observed low heritability. Future work could leverage high-quality 7T fMRI data to investigate whether this finding persists. As BEN holds great potential for understanding a wide range of neuropsychiatric diseases that are associated with aberrant complexity in certain brain regions and are often under genetic control [78], our findings on the heritability of BEN support that it may serve as an endophenotype for exploring the genetic etiologies of neuropsychiatric disorders. Further investigations are warranted to disentangle the common genetic basis between BEN and neuropsychiatric disorders.

### 4.2. Relationships between BEN and brain structure

In exploring the structural associations of BEN, we found that, in many cortical regions, BEN was phenotypically related to local cortical structure (especially SA and CM). Performing bivariate heritability analyses, we further revealed that specific genetic or environmental overlaps between BEN and cortical structure contributed to their phenotypic relationships. However, it should be noted that environmental overlaps encompassed correlated measurement errors and thus need to be cautiously interpreted [43]. Several studies using different anatomical characteristics or complexity measures indicated a significant association between signal complexity and brain structure. McDonough et al. [53] found that white matter integrity was positively associated with fine-scale multiscale entropy of fMRI signals but negatively associated with coarse-scale multiscale entropy. Liu et al. [54] adopted dispersion entropy to quantify the complexity of fMRI, revealing its association with cortical folding and myelination in the frontoparietal and attention networks. Our results extended the understanding of structure-complexity relationships by not only introducing new complexity measures and brain structural characteristics, but also by elucidating the link between structure and complexity at the level of genetic and environmental covariation. Our results improved knowledge about the neuroanatomical underpinning of fMRI temporal complexity and offered new insights into the relationship between structure and function.

Specifically, we observed significant negative correlations between BEN and SA in most cortical regions, suggesting that the enlargement of SA in these regions related to more predictable resting-state fMRI signals. It has been hypothesized that neural signals with low complexity may facilitate the establishment of phase relationships between distributed neural populations, leading to increased information communications across functionally segregated regions [20, 21, 22, 16]. Thus, brain signal patterns at low complexity may be instrumental for long-range functional integration. This can be supported by recent findings that lower complexity in resting-state fMRI signals was associated with stronger functional connectivity strength [22, 16] and better cognitive performance during language, working memory, and relational tasks [32]. Besides, the observed widespread negative correlations between BEN and SA implied that increased cortical surface area might promote long-range information communications across distributed cortical regions. In unimodal regions (particularly medial visual and somatomotor regions), we observed several positive correlations and many non-significant phenotypic correlations, indicating a lack of coupling between BEN and SA. This weak coupling pattern in unimodal regions may enable more flexible activity patterns in the medial visual and motor cortices upon the structural substrate, potentially promoting their capability to adapt to ever-changing external stimuli [79]. Upon further bivariate heritability analyses, we found that most of the phenotypic correlations were associated with a marked genetic correlation, suggesting a genetic overlap between BEN and SA. However, the genetic overlap was incomplete, with the observed genetic correlations being significantly different from 1 or -1. We also observed that substantial environmental correlations affected the phenotypic correlations between BEN and SA. Of note, environmental overlaps encompassed correlated measurement errors and thus need to be cautiously interpreted [43].

Moreover, positive phenotypic associations between BEN and CM were observed in our work. A recent hypothesis indicated that CM might inhibit synaptic plasticity [80]. Molecular evidence suggested that CM was detrimental to new axonal growth and synapse formation [80, 81, 82, 83]. It seems plausible that an increase in CM may impede information flow between separated cortical regions, therefore leading to positive associations between BEN and CM. Cortical regions with a significant correlation were primarily observed in high-order cognitive regions, including the prefrontal, lateral temporal, parietal, and cingulate cortices. Most of these regions have been reported to be lightly myelinated [80] and exhibit dramatic cortical expansion during evolution and development [84]. In addition, we found that the observed phenotypic correlations were mainly attributed to common environmental factors (including correlated measurement errors).

Only a few cortical regions exhibited significant correlations between BEN and CT. This observation is consistent with a recent study by Liu et al [54]. They investigated the complexity of fMRI via dispersion entropy and showed that cross-individual correlations between CT and fMRI complexity were narrowly distributed in the range of -0.2 to 0.2. With a predictive model, they showed that CT yielded very low predictions of individual complexity. Overall, our work and these findings collectively indicated a weak association between the complexity of fMRI signals and CT. Furthermore, the spatially sparse correlations between BEN and CT were partly associated with a significant environmental correlation (including correlated measurement errors).

### 4.3. Limitations

There are several limitations to our findings that need to be considered. First, the neuroimaging data of the young adult cohort that we used limited the generalizability of our findings in other age groups. Second, due to the dearth of accurate information regarding inter-individual shared environments in the HCP database, our analyses were performed in the AE model rather than the ACE model. This may lead to overestimates of genetic effects in our results. Third, the heritability analyses were limited by the fact that we evaluated the degree of inter-individual genetic similarity by kinship or pedigree data rather than genotype data in our models. Future studies can link genotype data to achieve a more precise picture of the level of genetic similarity between individuals.

## CRediT authorship contribution statement

Y.Zhen. and S.T. conceived of the study. Z.Z., S.T., H.Z., and Y.Zhen. designed the methodology. Z.Z., S.T., H.Z., X.W., L.L., Y.Y., Y.Zheng., Y.Zhen. contributed and implemented the investigation. Y.Zhen. and Y.Y. implemented the visualization. Y.Zhen., H.Z., S.T. and Y.Y. wrote and edited the manuscript.

## Acknowledgements

This work is supported by National Science and Technology Major Project (2022ZD0116800), Program of National Natural Science Foundation of China (62141605, 12201026), and Beijing Natural Science Foundation (Z230001). The HCP S1200 dataset was provided by the Human Connectome Project, WU-Minn Consortium (Principal Investigators: David Van Essen and Kamil Ugurbil; 1U54MH091657) funded by the 16 NIH Institutes and Centers that support the NIH Blueprint for Neuroscience Research; and by the McDonnell Center for Systems Neuroscience at Washington University.

## Data/code availability statement

The HCP S1200 dataset is publicly available at https://www.humanconnectome.org/study/hcp-young-adult/document/1200-subjects-data-release. The heritability analyses are performed through the Sequential Oligogenic Linkage Analysis Routines (SOLAR) software http://www.solar-eclipse-genetics.org and the R package solarius https://github.com/ugcd/solarius?tab=readme-ov-file.

## Supplementary information

The results of the robustness analysis are provided in Supplementary Materials.

