## Supplementary Materials for "The heritability and structural correlates of brain entropy"

Corresponding Author:

**Fig S1**. **A**. The heritability map of BEN, when the parameter setting was r=3, m=0.6$\sigma_{X}$. Only heritability estimates with $p<0.05/419$ were shown. **B**. Spatial correlation between heritability maps of different parameter settings: (m=2, r=0.5$\sigma_{X}$, main analysis) versus (m=3, r=0.6$\sigma_{X}$). The heritability estimates show very high consistency between the two parameter settings (Pearson correlation = 0.9913, uncorrected p < 0.0001). **C**. The phenotypic, genetic, and environmental correlations between local cortical structure (surface area (SA), cortical myelination (CM), cortical thickness (CT)) and regional BEN, when the parameter setting was r=3, m=0.6$\sigma_{X}$. For phenotypic correlations, only results with $p<0.05/1219$ were shown. For genetic and environmental correlations, only results with FDR corrected $p<0.05$ were shown. For each cortical structure, the top row denotes phenotypic correlations, the middle row denotes genetic correlations, and the bottom row denotes environmental correlations. We found no significant relationships between SV and BEN when the parameter setting was r=3, m=0.6$\sigma_{X}$.

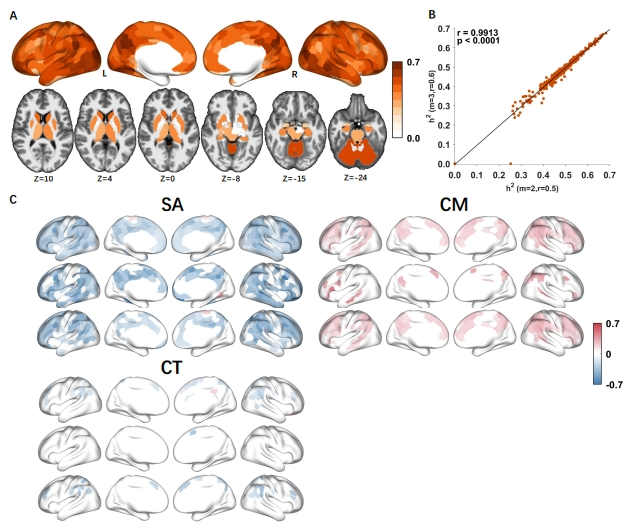

**Fig S2**. **A**. The heritability map of BEN, when performing global signal regression (GSR). Only heritability estimates with $p<0.05/419$ were shown. **B**. Spatial correlation between performing GSR and not performing GSR. The heritability estimates show high consistency (Pearson correlation = 0.9106, uncorrected p < 0.0001). **C**. The phenotypic, genetic, and environmental correlations between cortical structure (surface area (SA), cortical myelination (CM), cortical thickness (CT)) and regional BEN, when performing GSR. For phenotypic correlations, only results with $p<0.05/1219$ were shown. For genetic and environmental correlations, only results with FDR corrected $p<0.05$ were shown. For each cortical structure, the top row denotes phenotypic correlations, the middle row denotes genetic correlations, and the bottom row denotes environmental correlations. We found no significant relationships between SV and BEN when performing GSR.

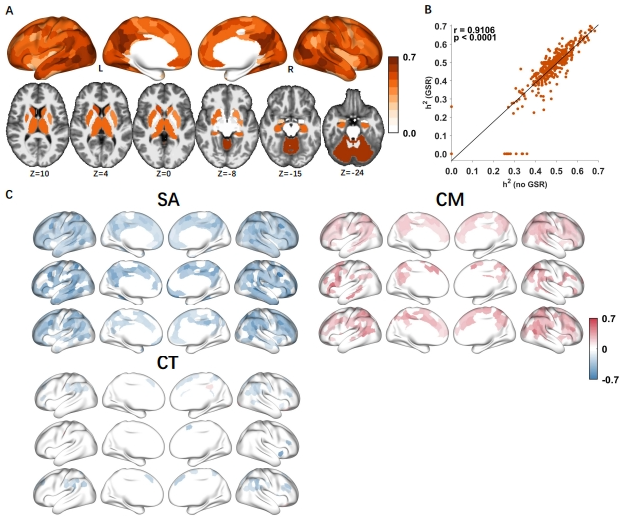
